# Origin of evolutionary bifurcation in an enzyme

**DOI:** 10.1101/2023.11.25.568631

**Authors:** Charlotte M. Miton, Eleanor C. Campbell, Joe A. Kaczmarski, Ferran Feixas, Adrian Romero-Rivera, Mahakaran Sandhu, Dave W. Anderson, Naoya Shatani, Sílvia Osuna, Colin J. Jackson, Nobuhiko Tokuriki

## Abstract

Evolution can lead to significantly distinct outcomes depending on the mutational path taken. Evolutionary bifurcation, in which two mutational trajectories segregate, becoming non-interchangeable over time, is the basis of diversification in all kingdoms of life. Here, we present a detailed molecular description of a bifurcation event that rapidly led to the emergence of two distinct enzymes from a common ancestor. When initiated from two starting points that differed by a single amino acid, the laboratory evolution of a phosphotriesterase (PTE) toward arylester hydrolysis resulted in different genetic and phenotypic outcomes. One trajectory led to a >35,000-fold increase in activity *via* the reorganization of its active site to achieve exquisite enzyme-substrate complementarity. The second trajectory gave rise to an evolved variant with a ∼500-fold increase in activity, but exhibiting an alternative substrate binding mode resulting from the destabilization of an active site loop. While initial mutations tend to dictate mutational accessibility, we rather observed the gradual divergence and specialisation of each trajectory, following the emergence of distinct molecular interaction networks. Intramolecular epistasis underlay pathway bifurcation by promoting unique synergistic interactions within each trajectory, while restricting the fixation of mutation across pathways. Our results illustrate how distinct molecular outcomes can radiate from a common protein ancestor and give rise to phenotypic diversity.

## Introduction

From higher organisms to molecules, life has evolved into “*endless forms most beautiful*” ^1^. This incredible diversity of evolutionary solutions arose from diverse factors, such as selection pressure, environmental influence, time, or even chance, which are inherent to evolutionary dynamics. For instance, an event as seemingly insignificant as a single mutation may dictate the path taken by evolution, leading to distinct evolutionary solutions. If so, evolution becomes highly contingent on a chance event. Evolutionary bifurcation, a phenomenon in which such a mutational event can trigger the segregation of two trajectories, giving rise to distinct and genetically incompatible species over time, has long fascinated biologists ^2^. The mechanisms underlying evolutionary bifurcation, following a chance event, remain largely unexplored, however.

Evolutionary bifurcation may stem from the complicated and intertwined nature of biological networks. Indeed, networks of amino acids, embedded within three-dimensional structures, underpin the remarkable efficiency of protein functions ^3,4^. While diverse residues shape the active site architecture to bind specific ligands, distal clusters of residues underlie protein motions and allostery *via* long-range coupling ^5^. Thus, functional evolution often entails the reorganization of the active site through the formation of novel interactions and the loss of existing ones; these alterations can trigger a signifcant rewiring of established intramolecular networks. Such rewiring is often accompanied by epistasis ^6^, whereby mutations interact in a non-additive fashion, a cardinal phenomenon that can dictate the fate of evolutionary trajectories ^7–9^. We hypothesize that bifurcation can arise when evolution follows alternative paths to rewire intramolecular networks, a consequence of extensive epistasis between mutations within, and across, evolutionary trajectories.

To explore the molecular underpinnings of evolutionary bifurcation in the laboratory, we conducted the parallel evolution of an enzyme, PTE phosphotriesterase, and observed the bifurcation of an ancestral lineage into two evolutionary pathways, from a common ancestor. Parallel experimental evolution, playing and replaying the “tape of life” ^10–13^, is a conventional approach to study the influence of chance on evolution. In fact, several studies already demonstrated that alternative trajectories can give rise to variants with distinct mutations ^14,15^. However, these studies have seldom elucidated the molecular changes that accompany the fixation of mutations, and it remains unclear how distinct solutions genetically segregate during evolution. Can a single mutation trigger a bifurcation by directly altering the effect of all dowmstream mutations, or is bifurcation rather a gradual phenomenon? By performing detailed genetic, biochemical, and biophysical characterizations of two evolutionary trajectories in unprecedented details (23 intermediates), we unveiled a molecular picture of an evolutionary bifurcation, which led to the emergence of two distinct molecular forms from a common ancestor. Our study illuminates how rapidly, but gradually, distinct molecular solutions can radiate from a common ancestor, giving rise to diversity.

## Results

### Laboratory evolution from neutral neighbours led to evolutionary bifurcation

We generated and compared two independent laboratory evolution experiments of PTE phosphotriesterase (PTE^WT^) toward the hydrolysis of an arylester, 2-naphthyl hexanoate (2NH) (**Fig 1a**). The reference trajectory, previously described in ref. ^16^, saw a 35,000-fold increase in arylesterase activity over 18 rounds of directed evolution from PTE^WT^ ^16^. H254R was the first mutation fixed along this trajectory (henceforth referred as ‘*Arg* trajectory’); it increased the arylesterase activity by ∼3.5-fold and became a key residue, as other mutations accumulated (**Fig 1b**) ^16–18^. The prominent role of H254R led us to hypothesize that distinct amino acids at position 254 could set the stage for a bifurcation event and the creation of alternative adaptive pathways (**Fig 1c**). A mutational scanning at position 254 in PTE^WT^, *i.e.*, mutating His254 to all 19 amino acids, revealed that most substitutions are mildly deleterious or neutral. We chose Ser254, a residue with a different charge and size from arginine, and nearly neutral with respect to the arylesterase activity in PTE^WT^ (1.2-fold increase in 2NH hydrolysis) (**Fig. 1b)**. We then performed directed evolution to enhance PTE’s arylesterase activity, under identical experimental conditions (‘*Ser* trajectory’, **Fig. 1b**). Note that PTE variants are named according to the evolutionary round and trajectory in which they occurred: for example, R8 is the round 8 variant fixed along the *Arg* trajectory, while S8 is the corresponding variant in the *Ser* trajectory.

**FIGURE 1.**
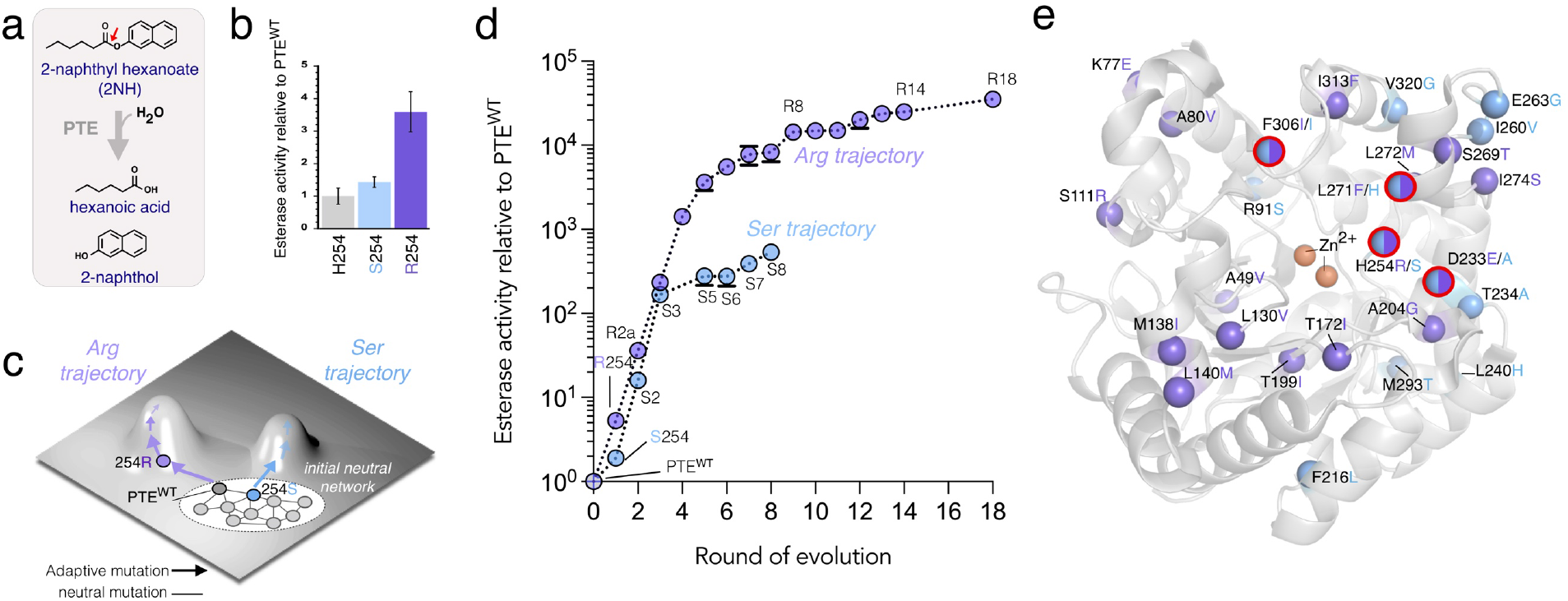
Phenotypic and genotypic adaptation along the *Ser* and *Arg* trajectories. **a**, Reaction scheme of 2-naphthyl hexanoate hydrolysis by PTE. **b**, Esterase activity of PTE^WT^, PTE^S1^ (H254S) and PTE^R1^ (H254R) in cell lysate normalized on PTE^WT^. **c**, Schematic representation of the evolutionary trajectories created in this study. Directed evolution was performed from two genetic neighbours, PTE^WT^ (His254) and PTE^S1^(Ser254). **d**, Changes in esterase activity during directed evolution, measured in cell lysate and normalized on PTE^WT^. Error bars represent s.d. of duplicate measurements from three independent experiments. **e**, Mapping of the mutations fixed during directed evolution on the PTE^WT^ structure (PDB: 4pcp), represented as spheres. Red circles highlight positions mutated in both trajectories. Two active-site zinc ions are shown as golden spheres.

Contrary to *Arg*, our efforts to evolve highly active variants along the *Ser* trajectory were thwarted (∼500-fold in PTE^S8^ *vs.* >35,000-fold in PTE^R18^, **Fig. 1d**). Despite initially differing by a single amino acid, the two bifurcated trajectories reached genetically distinct solutions: all mutations but F306I are unique to their respective lineages (**Fig. 1e**). Overall, the *Arg* mutations are scattered across the whole structure, particularly within Loop 4-8, whereas the *Ser* mutations are restricted to Loop 6-8. In addition to His254 and Phe306, Asp233 and Leu271 are also mutated in both trajectories, albeit to different amino acids. While D233E / L271F arose with H254R in the *Arg* trajectory, D233A / L271H were fixed along H254S in the *Ser* trajectory. Interestingly, these positions all surround the active site, and can - depending on the amino acid present - form interactions with the substrate and with one another (**Fig. 1e**). We refer to the ‘REF’(Arg254-Glu233-Phe271) and the ‘SAH’ (Ser254-Ala233-His271) triplets in each trajectory hereafter.

### The rewiring of intramolecular networks created alternative structural solutions

Next, we sought to investigate the molecular changes underlying each trajectory. We first obtained 10 crystal structures of all seven *Ser* variants, including PTE^S5^ in complex with a transition state analogue (HLN, hexylphosphonate 2-naphthyl ester). We then compared them to 16 structures of 13 variants from the *Arg* trajectory, including PTE^R18^ (**Fig. 2a-d**). Moreover, we performed molecular dynamics (MD) simulations (**Fig. 2e,f**) and ensemble refinement (ER) (**Fig. 2g**) on 6 and 16 structures, respectively, to investigate potential differences in structural motions along the two trajectories. We previously demonstrated that along the *Arg* trajectory, the active site pocket was reshaped to provide exquisite enzyme-substrate complementarity, supporting the high catalytic efficiency of PTE^R18^ (**Fig. 2a**) ^16^. Here, we found that HLN – and by extension 2NH – bind in strikingly different orientations in PTE^S5^ (**Fig. 2b**). While the hexyl chain adopts a similar position in both enzymes, the naphthyl moiety of the substrate exhibits a kinked conformation in PTE^S5^. This orientation, albeit productive, appears suboptimal and more solvent-exposed, consistent with a ∼130-fold lower activity in PTE^S5^ compared to PTE^R18^.

**FIGURE 2.**
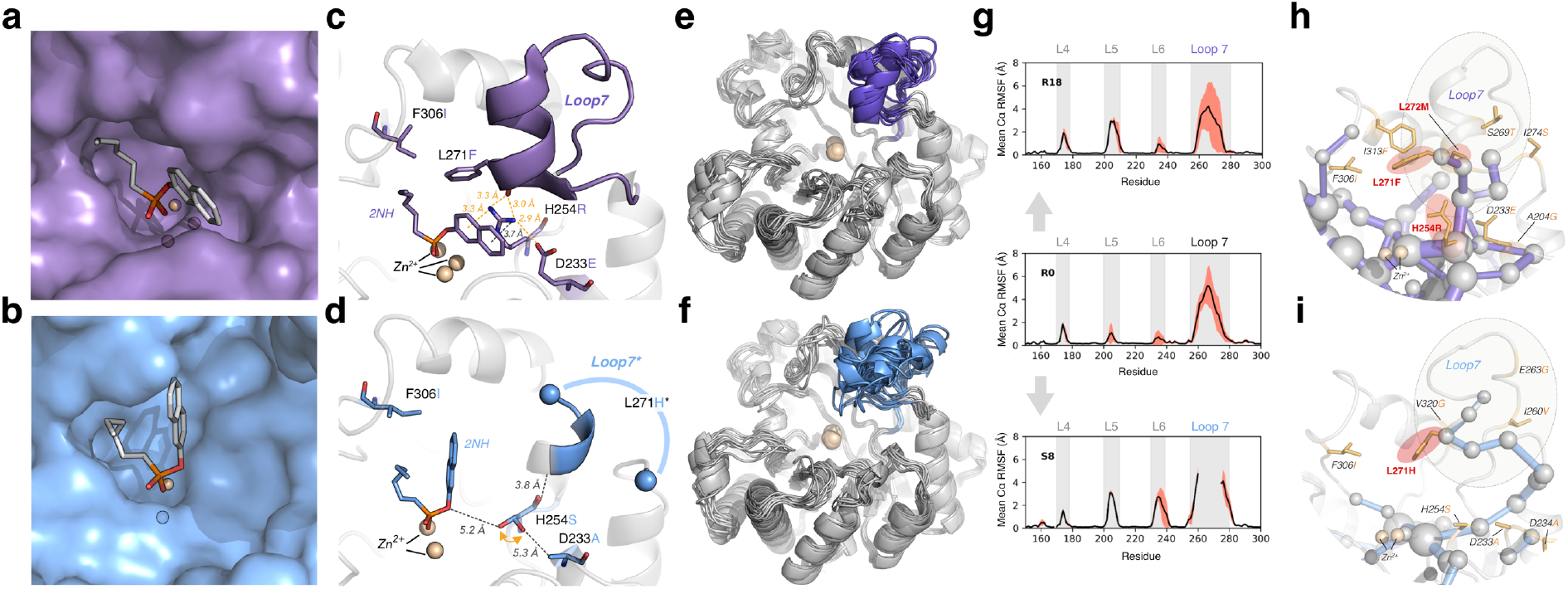
A new structural adaptation in the *Ser* trajectory reveals an alternative molecular solution for ester hydrolysis. **a,b,** Surface representation of the crystal structures of (**a**) PTE^R18^ (PDB 4e3t) and (**b**) PTE^S5^ (PDB 6B2F) with bound transition state analogue, HLN shown as sticks**. c**, A novel hydrogen bonding network between residues 233-254-271 in PTE^R18^ enhances leaving group stabilisation by π-cation interactions and stabilizes the closed conformation of loop 7. **d,** In PTE^S8^ (PDB 6AML), adaption disrupted the 233-254-271 network, resulting in substantial substrate repositioning and increased Loop7 dynamics (too disordered to be modelled, thus represented as a blue line). HLN is overlaid from PDB 6B2F. The substrate adopts an alternative binding mode in PTE^S8^, with suboptimal enzyme-substrate complementarity. Orange arrows emphasise dual residue conformations. Atomic distances are indicated as dotted lines: orange if ∼3 Å, or black if ≥ 3.5 Å. **e,f**, Molecular dynamics simulations of (**e**) PTE^R18^ and (**f)** PTE^S8^ illustrating significant changes in Loop 7 (colored) dynamics. **g,** Mean root mean square fluctuation (r.m.s.f) of the Cα backbone of PTE^R18^, PTE^WT^ and PTE^S8^, revealed by ensemble refinement. The single line represents the mean r.m.s.f across five replicate refinements with uncertainty (s.d.) shaded in red. **h,i,** Shortest Path Map (SPM) analysis of (**h**) PTE^R18^ and (**i**) PTE^S8^ demonstrates the evolution of distinct intramolecular networks traversing the PTE active site and connecting with Loop 7. The size of the edges (purple or blue lines, respectively) and spheres (grey) represent the strength of the network, *i.e.*, the relevance of a given position and the degree of connectivity between residues. Mutations are shown as orange sticks; red halos represent trajectory mutations directly on the shortest path, indicative of a correlation between evolutionary optimization and correlated movements. Zinc ions are shown as golden spheres; they illustrate the location of the catalytic centre. **See Ext. Fig. 2** for full view.

These two binding modes seem to originate from contrasting structural adaptations, resulting from the establishment of distinct intramolecular networks (**Fig. 2c,d**). In the *Arg* trajectory, a hydrogen-bonding network arose between active site residues 254-233-271(**Fig. 2c)**. As previously shown, Arg254 adopted a bent conformer (Cβ-Cγ-Cδ-Nε dihedral angle ∼40−70°) enabling the productive binding of 2NH observed in PTE^R18^ by forming cation-π interactions with the naphthyl moiety of the substrate. The Arg254 bent conformer is directly stabilized by hydrogen-bonding interactions with D233E and Loop 7 residues, such as F271L ^16,18^. Consequently, Loop 7 motions are greatly reduced in PTE^R18^ (**Fig. 2e,g)** ^18^. On the contrary, PTE^S5^ mutations do not induce the formation of hydrogen-bonding interactions between positions 233-254-271, resulting in a dramatically destabilized Loop7 in PTE^S3-S8^ (**Fig. 2d, f, g**), such that Loop 7 residues are too mobile to be observed in the electron-density map. Furthermore, we computed shortest path maps (SPM), an approach that identifies networks of residues with coupled motions, *i.e.*, communication pathways, during MD simulations ^19,20^. We found that distinct intramolecular networks were formed along each trajectory (**Fig. 2h,i**). In PTE^R18^, the transduction of structural motions seems to span various active site loops. In particular, we observed a high number of pathways connecting Arg254 to Loop 7, where high residue connectivity could help minimize Loop 7 dynamics to enhance catalysis (**Fig. 2h**). By contrast, in PTE^S8^, communication pathways are atrophied and overall, disconnected, which suggests that the intramolecular network mediating Loop 7 dynamics is significantly distinct in this enzyme (**Fig. 2i**). Taken together, these results suggest that, despite an identical selection pressure, two bifurcated trajectories led two distinct molecular solutions, *i.e*., exhibiting distinct intramolecular networks, structural dynamics, and perhaps more surprisingly, enzyme-substrate complexes.

### Differential mutational tolerance supports distinct adaptive mechanisms and exposes epistasis

Exploring mutational tolerance can inform on the contribution of a given residue to the enzymatic function. To this end, we mutated each key position 254, 233, 271 and 306 to all 19 amino acids in PTE^WT^, PTE^S5,^ and PTE^R18^. In PTE^R18^, all R254X substitutions decreased the activity by >1,000-fold, including serine and histidine. Similarly, mutations at position 271 and 306 are largely deleterious, albeit to a lesser extent. Mutations at 233 are only marginally deleterious in PTE^R18^. We observed contrasting mutational reponses in PTE^S5^: mutations at position 306 are the most deleterious, whereas 254 mutations exhibit a binary behaviors, *i.e.*, they either appear neutral or deleterious. Mutations at 233 are mildly deleterious and almost completely neutral at position 271. Differential mutational responses in the two evolved variants are consistent with structural observations: disrupting any element of the 233-254-271 network supporting the Arg254 - substrate interaction is strongly detrimental along the *Arg* trajectory. The corresponding mutations within the SAH triplet (*Ser* trajectory), which no longer forms an interaction network, are thus less disruptive. Note that I306X mutations are poorly tolerated in both variants, confirming that F306I is a key mutation in both trajectories, where it mediates the binding of the alkyl side-chain.

The saturation libraries at 233-254-271 contain ‘back to wild-type’ reversions, *e.g.*, R254H or S254H, and ‘swapping’ mutations between the *Ser* and *Arg* trajectories, *e.g.*, R254S or E233A. A detailed characterization of these mutations can expose if and how each lineage has specialized to utilize a given amino acid. As expected, most reversions are deleterious in both enzymes, especially in PTE^R18^, which demonstrated the relevance of these positions for the evolved function, and confirmed that these lineages have significantly diverged from PTE^WT^ (**Fig. 3a**). Swapping alleles between *Arg* and *Ser* trajectories at position 233 (Glu *vs*. Ala), 254 (His *vs.* Arg), and 271 (Phe *vs*. His), individually or altogether (REF *vs*. SAH), demonstrated that hybrid variants are less fit than their parents (**Fig. 3b**). For example, R254S in PTE^R18^ decreased the arylesterase activity by ∼430-fold; S254R while being deleterious in PTE^S5^ or PTE^S8^, is limited to a 2-fold decrease. This mutual incompatibility between amino acids at position 233-254-271, a phenomenon reflecting reciprocal-sign epistasis, suggests that each residue in the network plays a specialized role in their respective evolved variant, rendering them non-functional in another background. The X-ray structures of these two ‘swapped’ variants provided molecular insights into this incompatibility: PTE^R18^+R254S exhibits a distorted active site architecture, including high flexibility in Loop 7. On the other hand, introducing S254R into PTE^S5^ has little functional or structural effect: the 1.5-fold decrease in arylesterase may be attributed to a slight steric clash with the substrate bound in the *Ser* conformation (*i.e.*, as in PTE^S5^ + HLN structure).

**FIGURE 3.**
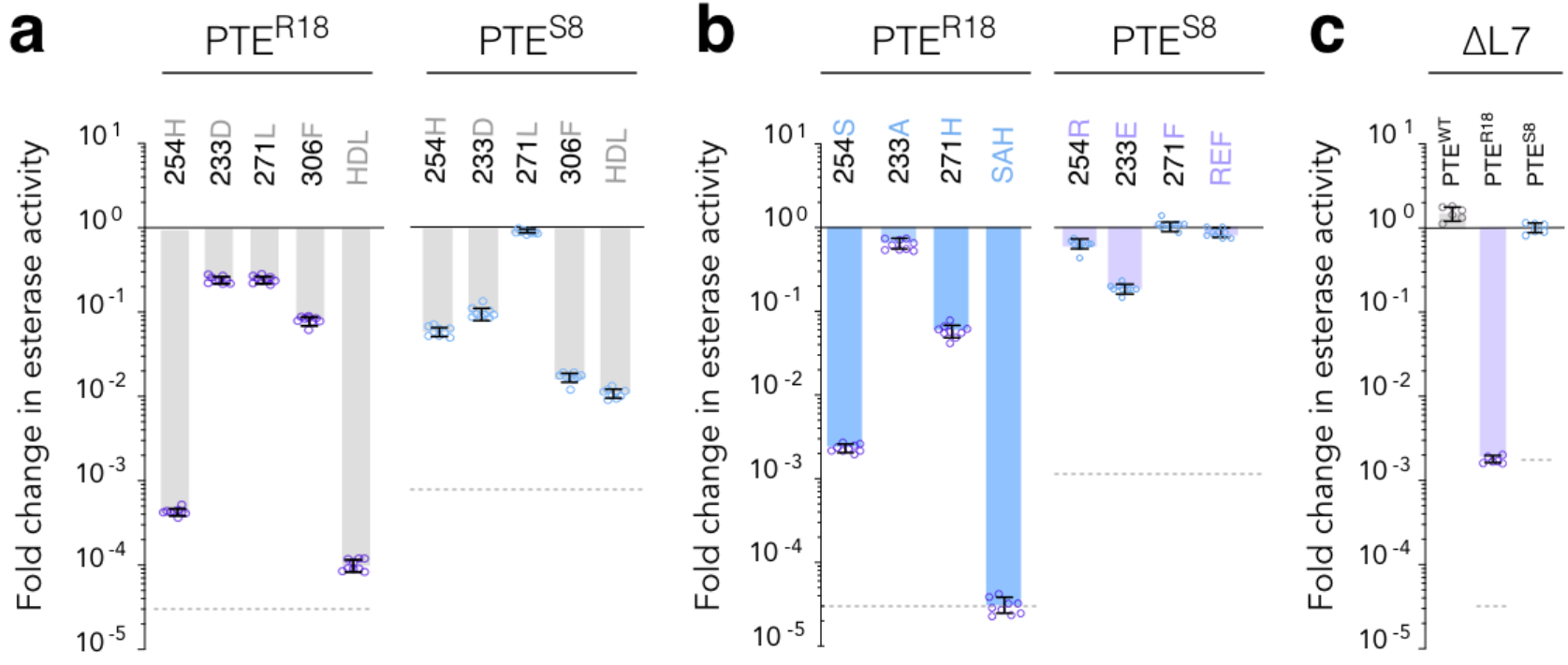
Differential tolerance to mutations exposes distinct molecular mechanisms and epistasis. **a-c,** Fold change in esterase activity upon mutating positions 254, 233, 271, and 306, individually or in combination, in PTE^R18^ and PTE^S8^, normalized on the corresponding background. **a,** Effect of ‘back-to-wt’ mutations (H, D, L, F, HDLF), *i.e.*, reversion to the PTE^WT^ genotype. **b,** Effect of ‘swapping’ mutations, *i.e.*, introducing the PTE^S8^ mutations (S, A, H, SAH) into PTE^R18^, and reciprocally (R, E, F, REF). **c,** Effect of deleting Loop 7 in PTE^WT^, PTE^S8^ and PTE^R18^. All experiments were measured in duplicate over three independent experiments in cell lysate. Error bars represent s.d. from the mean. The dotted lines represent the maximum fold difference, *i.e.*, when a given mutation decreases the function down to the PTE^WT^ level, cancelling the beneficial effects of all other mutations fixed in the PTE^R18^ or PTE^S8^ backgrounds.

Finally, we probed the role of Loop 7 along each trajectory by constructing Loop 7 deletion variants (named ΔL7) on the PTE^WT^, PTE^S5^ and PTE^R18^ and obtaining their crystal structures (**Fig 3c**). While ΔL7 does not prevent soluble expression in any variants, the functional and structural consequences differ substantially. ΔL7 has virtually no effect on the arylesterase activity, nor on the active site integrity of PTE^WT^ and PTE^S8^. By contrast, ΔL7 is extremely detrimental to PTE^R18^; the activity decreased by ∼1,000-fold. We found that the active site organization became very disrupted, particularly in the vicinity of Arg254: this key residue now exhibits, once again, two conformers (bent and extended), which likely compromised catalysis. Taken together, these results confirm the existence of extensive underlying epistasis, which could have arised from potential alterations of pre-existing interaction networks.

### The bifurcation of the *Ser* and *Arg* trajectories is gradual

We established the relevance of studying 233-254-271 in the evolved variants. Yet, several other mutations (15 in *Arg* and 9 in *Ser* trajectories, respectively) were also accumulated in the intermediate variants along each path. To acquire a time-resolved picture of evolutionary bifurcation, we introduced “254 reversions” (S/R254H**, not shown**), “254 swaps” (R254S, S254R, **Fig. 4a**), and ΔL7 (**Fig. 4b)** mutations at various stages along the trajectories. We found that the effect of every mutation was gradually altered over the course of evolution. For instance, in the *Arg* variants, R254S, R254H, and ΔL7 only initially decreased the arylesterase activity by a few fold; these effects gradually and significantly increased in later rounds, up to >1,000-fold. While the effects in the *Ser* trajectory appear less drastic, the initial beneficial effect of S254R and the deleterious effect of ΔL7 gradually disappear, becoming neutral as evolution unfolds. This remarkable graduality in mutational effects is a manifestation of extensive epistasis: most, if not all, mutations interact with position 254 and/or Loop 7 and alter the functional role of these two key elements. Furthermore, these observations suggest a gradual and extensive rewiring of the original intramolecular networks that existed in PTE^WT^ during evolution. This rewiring is distinct along the two trajectories, *Arg* and *Ser* adapted to their initial mutation differently: Arg254 gradually became essential to the function in the *Arg* trajectory, whereas Ser254 (and position 254 overall) became largely insensitive in the *Ser* variants.

**FIGURE 4.**
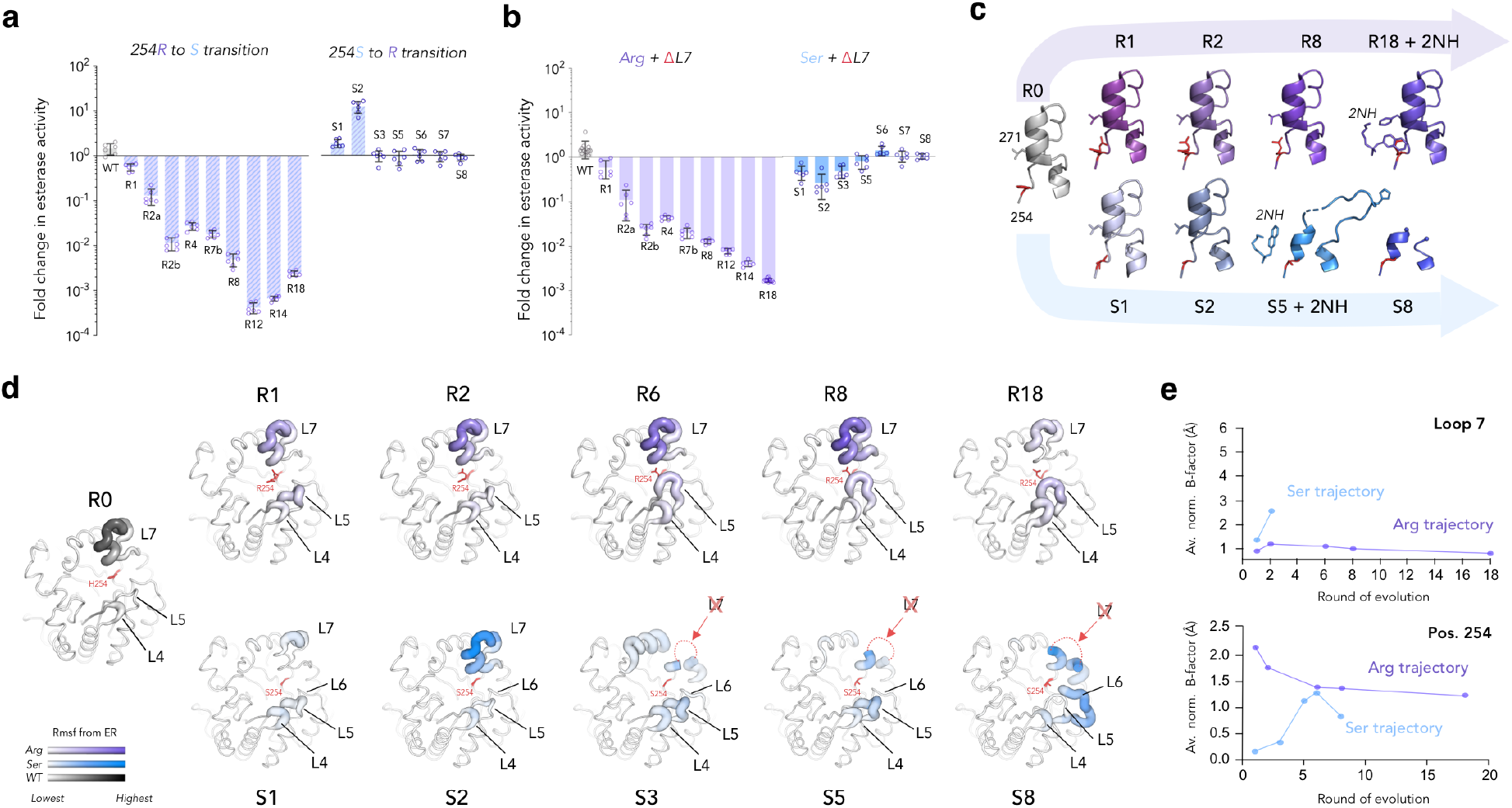
Distinct adaptive mechanisms gradually evolved and lead to altered Loop7 dynamics along the *Ser* and *Arg* trajectories. **a,** Effect of swapping mutations at positions 254 along the *Ser* and *Arg* trajectories. **b,** Effect of deleting Loop 7 (ΔL7) along the *Ser* and *Arg* trajectories. PTE^WT^-ΔL7 is shown in grey. **c,** Evolution of Loop 7 structure along the *Ser* and *Arg* trajectories. The density of Loop 7 is missing in PTE^S8^ due to high mobility. **d,** Ensemble refinement of key variants along the *Ser* and *Arg* trajectories, respectively. R.m.s.f are displayed on the structure as colored putty, where gradient and thickness, using a scaling factor, reflect the extent of structural motions. The residue at pos. 254 are shown as red sticks. **e,** Evolution of averaged baseline-normalized B-factors along the *Ser* and *Arg* trajectories. The changes in structural motions along the trajectory varies for different regions. (*Top panel*) Loop 7 peaked at round 2 along the *Arg* trajectory and decreased continuously until round 18. In the *Ser* trajectory, it is presumed to rich a high B-factor from round 3 onwards. (*Bottom panel*) In both trajectories, pos.254 peaked in round 1 and was stabilized by round 6.

Structural, ER and B-factor analyses of all variants provided evidence for a gradual network rewiring (**Fig. 4c-e and Ext. Fig 6**). We found that the structural changes strikingly mirror the functional ones. First, we observed a unique shift in Loop 7 dynamics along the *Ser* trajectory, with a significant increase in Loop 7 dynamics throughout evolution (**Fig. 4c)**. The extent of the conformational fluctuations sampled in PTE^S5^ can be visualized in the liganded structure with HLN, where Loop 7 became visible again (but not in the apo structure) (**Fig. 4c)**. Furthermore, we showed that the rmsf (ER) and B-factors of Arg254 gradually decreased along the *Arg* trajectory, as more mutations accumulated (**Fig. 4d,e**). In parallel, Loop 7 was initially destabilized following the fixation of H254R (round 1), but became gradually stabilized over the course of evolution. By contrast, Loop 7 is increasingly - and gradually - destabilized along the *Ser* trajectory (**Fig. 4d,e**), in correlation with our functional observations (**Fig. 4b**). Our results reveal the molecular mechanisms by which the gradual functional optimization of PTE was accompanied by a gradual evolutionary optimization of the structural motions of Loop 7 and Arg254.

### Epistasis causes the divergence of protein lineages across the fitness landscape

Last, we sought to identify specific epistatic interactions causing the bifurcation of the two trajectories. To this end, we generated and characterized the local fitness landscape of the key triplet positions, 254 (His, Ser and Arg), 233 (Asp, Ala and Glu) and 271 (Leu, His and Phe), by constructing all 27 combinatorial genotypes on the background of PTE^WT^+F306I (**Fig. 5a**). Interestingly, the triplets accumulated in each trajectory (REF and SAH) exhibit distinct epistatic trends. The *Arg* trajectory exhibited strong pairwise positive epistasis, *e.g.*, H254R-D233E, and H254R-L271F (**Fig. 5a,b**). This suggests that the initial H254R mutation enabled the fixation of subsequent mutations D233E and L271F, paving the way for the *Arg* trajectory to evolve. It may also explain the stepwise creation of the 254-233-271 network, each position acting as a building block in this interaction network that stabilizes the substrate and the active site. By contrast, the *Ser* trajectory comprises more negative epistasis, whereby double mutants exhibit lesser effects than the additive effect of the individual mutations (**Fig. 5b)**. This negative epistasis may illuminate how the *Ser* trajectory relied on the disruption of the 233-254-271 network: each mutation damages the network in a similar, but not additive, manner. Beyond the trends observed within each trajectory, we identified several examples where mutation(s) in one trajectory restricted the appearance of mutations in the other, due to sign epistasis. For instance, H254S (*Ser*) diminished the effect of D233E (*Arg*), and concomitantly H254R (*Arg*) that of D233A (*Ser*) (**Fig. 5c**). Furthermore, when D233A (*Ser*) and L271F (*Arg*) are fixed after H254R (*Arg*), they exhibit reciprocal sign epistasis causing a 66-fold lower arylesterase activity than their additive effect (**Fig. 5c**). H254R (*Arg*) and L271H (*Ser*) also exhibit sign epistasis, which implies that the fixation of H254R in the *Arg* trajectory restricted the emergence of L271H in this trajectory (**Fig. 5c**). These results are consistent with the fact that *Arg*/*Ser* hybrids are incompatible with one another (**Fig. 3**) and explain, at the molecular level, the bifurcation of these lineages. Taken together, prevalent epistasis facilitated the opening of paths to mutations in their own trajectory, while restricting others, resulting in highly incompatible and distinct evolutionary outcomes.

**FIGURE 5.**
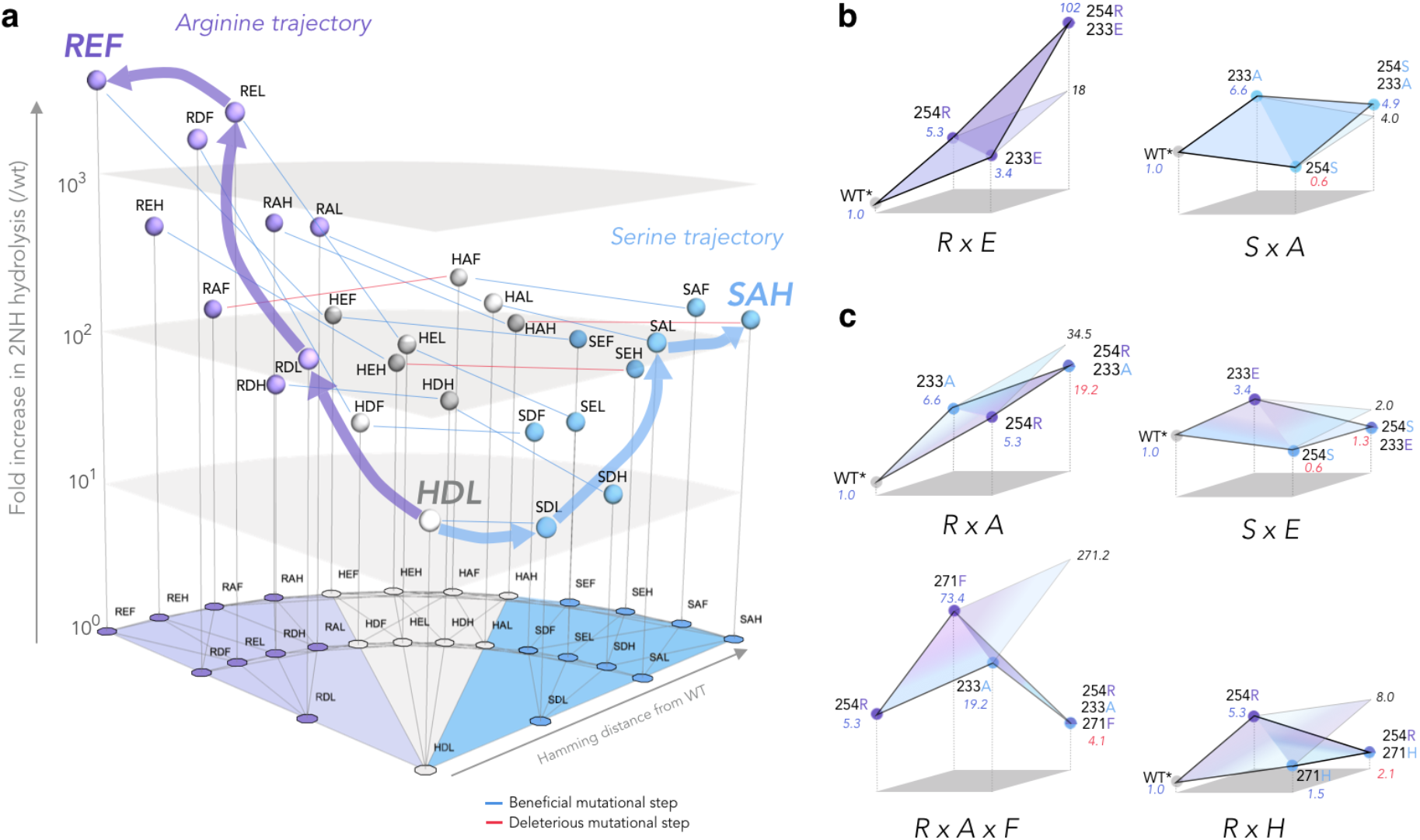
Epistasis drives the divergence of the *Ser* and *Arg* trajectories across the fitness landscape. **a**, Fold increase in 2NH activity of all 27 genotypes combining mutations at pos. 254-233-271 on the background of PTE^WT^ + F306I (called HDL*). At the basis of the landscape, nodes (purple: R254; grey: H254 and blue: S254) represent genotypes (254X, 233X, 271X) and edges (grey lines) represent single mutational steps between neighbours. The arylesterase activity of the 27 variants is normalized on the activity of PTE^WT^. Purple and blue arrows represent the trajectories selected during directed evolution from HDL* to REF (*Arg* trajectory) or to SAH (*Ser* trajectory), respectively. At first, the sub-optimal *Ser* trajectory can still access the optimal *Arg* trajectory (dark blue edges), *via His* intermediates. They gradually become separated by a deleterious valley were fitter genotypes are no longer accesible (red edges). Grey surfaces indicate logarithmic scale of catalytic activity. **b-c**, Unique epistatic effects occurring along both lineages explain the selection of distinct alleles at position 233 and 271. **b** Effect of mutation D233E *vs.* D233A when combined with H254R or H254S. **c**, Genetic incompatibility results from negative epistasis when D233A occurs on the background of H254R and L271F. Similarly, mutation L271H masks the beneficial effect of H254R, due to negative epistasis.

## Discussion

In this study, we presented a detailed molecular picture of evolutionary bifurcation, where two distinct and genetically incompatible enzymes rapidly evolve from a common ancestor. Beyond this case study, our observations have broader implications for long-sought questions in protein evolution and engineering. For instance, how does protein evolution lead to different local maxima? Our study shows, for the first time, that genetically and molecularly distinct enzyme solutions exist in the fitness landscape, a stone’s throw away from a common ancestral state, and from one another. While we found that the initial mutations (H254R *vs.* H254S) triggered the bifurcation by promoting and restricting access to distinct subsets of mutations, the cumulative impact of subsequent mutations was essential to achieve pathway segregation. As more mutations accumulate, they gradually rewire interactions, expand or annihilate existing networks, reinforcing their own role within these networks. Our results, consistent with previous studies ^18,19^, further suggest that intertwined networks could reflect the fine-tuning and optimization of structural dynamics along the evolutionary trajectories of highly efficient enzymes. Such sophisticated coupling did not occur along the *Ser* trajectory, which also explains its dimisnished catalytic efficiency.

Concomittantly, we demonstrate that newly evolved networks can become incompatible with residues contributing to distinct networks, resulting in the creation of two uniquely distinct “molecular species”. While we report on a single enzyme, such mechanisms are likely relevant to other biological systems, where substantial networks (*e.g.*, intra- or inter-molecular, and genetic networks) underlie the function. In this study, we convincingly demonstrate that, while the fixation of key initial mutations may set the stage for network rewiring, it is the accumulation of consecutive mutations, and how they interact with the initial mutations, that will cause evolutionary diversification. Such snowball effects can lead to the rapid bifurcation of evolutionary pathways, and ultimately, the adaptive radiation of species ^21,22^.

A question arises regarding the frequency of such “bifurcating mutations” within the protein sequence space. Our study might constitute an exceptional case whereby mutations at the same position led to a clear bifurcation and diversification. Indeed, position 254 is located at the heart of the active site and can form direct interactions with the substrate. Other positions in PTE are unlikely to cause similar effects. Furthermore, it is generally assumed that only a relatively small fraction of mutational combinations will give rise to epistasis in a protein, let alone sign epistasis, an even rarer phenomenon. Moreover, the magnitude of epistasis is rarely as drastic. Therefore, only a handful of mutational pairs may be able to trigger evolutionary bifurcation. Yet, our study illustrates that, once such pairs are identified, rapid and drastic diversification can unfold.

Nonetheless, when we consider long-term evolutionary time scales, evolutionary bifurcation caused by chance, *i.e.*, seemingly insignificant mutational events, can be substantial. During genetic divergence, *i.e.*, neutral drift, multiple mutations will accumulate, while maintaining the primary function of a protein. Since genetic divergence among protein orthologs can easily exceed >100 amino acid changes, these functionaly neutral mutations can set the foundation for evolutionary bifurcation by dictating potential rewiring strategies. Indeed, several recent studies showed that experimental evolution from homologous sequences resulted in significantly different evolutionary outcomes ^23,24^. Thus, protein evolution is highly unpredictable, unless we understand the molecular details, that support protein functionality, including the mechanisms underlying the emergence and functionaing of intricated interaction networks in proteins.

## MATERIAL AND METHODS

### Methods Summary

Libraries of *Ser* variants were created by error-prone PCR or DNA shuffling and cloned into a pET-Strep vector, as described in ref. (Tokuriki et al., 2012). Variants were expressed in *E. coli* BL21 (DE3) in the presence of GroEL/ES chaperones (pGro7 plasmid, TAKARA Bio, USA). After a pre-screen on agar plates, active variants were expressed in 96-well plates and assayed for improved 2NH hydrolysis in cell lysate. To determine the effect of mutations in different backgrounds, additional variants were constructed by site-directed mutagenesis and assayed in cell lysate. All variants were also purified by Strep-tactin chromatography and initial rates were measured at a fixed concentration of 2NH. Michaelis Menten parameters were determined for a subset of variants and all *Ser* variants were crystallized and their structures were solved.

### Material

2NH was originally purchased from Sigma; it was later kindly gifted to us by the Tawfik Lab. Fast Red TR hemi (zinc chloride) salt (Fast Red) was purchased from Sigma. The TS analogue of 2NH (HLN, hexyl(naphthalen-2-yloxy)phosphinic acid) hexylphosphonate 2-naphthyl ester was synthesized by the Tawfik lab based on previously published procedures ^16,25^.

### Methods

#### Cloning

The gene encoding PTE^WT^ (GenBank accession: KJ680379) is identical to the one used in refs. ^16,18,26^. Note that PTE^WT^ (Uniprot accession: A0A060GYS1) had previously been evolved for improved expression levels in *E. coli* ^27^, and contains seven amino acid mutations (I106L, F132L, K185R, D208G, R319S, A176V, V341I) relative to the naturally occurring PTE (Uniprot accession: P0A434). All PTE *Ser* variants were cloned with NcoI/XhoI sites into a kanamycin-resistant pET-27-STREP vector for evolution and purification. These constructs differ from the *Arg* variants (Ampicillin, NcoI/HindIII) to avoid cross-contamination with the original trajectory. PCR products and vectors were digested with FastDigest restriction enzymes (Thermo Scientific) for 1-2H at 37 °C and subsequently purified from a 1% agarose gel using a DNA micro-elute purification Kit (OMEGA). Ligations were performed at a vector: insert mass ratio of 3:1 using T4 DNA ligase (Thermo Scientific) at 16 °C overnight. Prior to transformation, reactions were purified using the DNA micro-elute purification Kit (OMEGA). Transformation into electrocompetent *E. cloni* 10G (Lucigen, Middleton, WI, United States) reliably yielded >20, 000 colonies from 50-100 ng of DNA per ligation. ACP (acyl phosphatase) cloned into the same pET-27-STREP vector served as a negative control for 2NH activity measurements. All constructs were validated by Sanger DNA sequencing (GENEWIZ).

#### Single point mutations

Mutants were constructed by: 1) site-directed mutagenesis as described in the QuickChange Site-Directed Mutagenesis manual (Agilent), or 2) using type-II restriction cloning, adapted from Golden Gate cloning ^28^. Briefly, after re-cloning PTE variants into a pET-27-ΔLguI_STREP vector (without LguI/BsaI sites in the backbone), whole plasmid amplification was carried out by PCR with primers containing the LguI or BsaI sites, as well as the desired mutation. The amplified DNA was purified using a DNA micro-elute purification Kit (OMEGA), treated with DpnI and LguI or BsaI (Fastdigest, Thermo Scientific) for 2-3H at 37 °C. DNA was further purified with the micro-elute Kit, self-ligated (<50 ng) overnight at 16 °C and transformed into *E. coli* cells, as described above. All mutations were validated by Sanger DNA sequencing.

#### Directed evolution libraries creation

*epPCR libraries* - PTE libraries were generated by random mutagenesis with error-prone PCR (epPCR) using the ‘wobble’ base analogues dPTP and 8-oxo-dGTP (TriLink), with slight alterations from reference ^16^. The first out of two PCR reactions was performed over 20 cycles (2 min at 95 °C; 20x: [20s at 98 °C, 15s at 58 °C, 45s at 72 °C]; 2min at 72 °C). The 50 μL reaction contained 1ng of plasmid template, 0.3 µM primers, 0.75 µM dPTP or 60 µM 8-oxo-dGTP, 0.25 mM of each of the four standard dNTPs, 2.5 mM of MgCl_2_ in the supplied buffer. The PCR products were purified with the DNA micro-elute purification Kit (OMEGA), treated with DpnI for 2H at 37 °C to digest the template plasmid, purified again, and quantified with a Nanodrop (ThermoScientific). DNA from each epPCR was then combined in equimolar quantities and 10 ng served as a template for the second reaction. The second PCR uses the same protocol as above, except with nested primers and the use of regular dNTPs. The mutated inserts obtained were then digested and cloned into pET-27-STREP as described above. This method resulted in the creation of libraries that carried an average of 1.5 amino-acid mutation/gene with high reproducibility, calculated from 10 randomly selected clones after transformation into *E. cloni* and prior screening. Shuffling mutagenesis – At rounds 3 and 5, StEP PCR (staggered extension protocol) ^29^ was used to recombine positive variants isolated from epPCR libraries. A 100 µL reaction with 500 ng of DNA template, which consisted of equimolar amounts of DNA from each variant of interest, was shuffled in the PCR reaction. The first PCR conditions are identical to the second epPCR, except for the primers used at 0.1-1nM and the PCR cycle modified as follows: 100 cycles (5 min at 95 °C; 100x: [30s at 95 °C, 5s at 58 °C]). PCR products were digested with DpnI, purified with the micro-elute kit and 1.5 µL were used as a template for a second PCR: 20 cycles (2 min at 95 °C; 20x: [20s at 98 °C, 15s at 58 °C, 45s at 72 °C]; 2min at 72 °C) using the EconoTaq Plus protocol (Lucigen, Middleton, WI, United States). The PCR products were then digested, purified from DNA electrophoresis gel and cloned into pET-27-STREP as described above.

#### Saturation libraries creation

Libraries of PTE variants containing all 20 amino acids at position 254X, 233X, 271X and 306X for PTE^WT^, PTE^R18^ and PTE^S8^ were independently created with primers containing a degenerated ‘NNS’ codon at these positions, followed by type II restriction cloning as described above.

#### Arylesterase assay on agar plates

Plasmids were extracted from E. cloni cells using a QuiaPrep mini spin kit (Quiagen) and re-transformed into *E. coli* BL21 (DE3) containing a pGro7 plasmid for overexpression of the GroEL/ES chaperone system (Takara Bio, Shiga, Japan). ACP (acyl phosphatase) cloned into the same pET-27-STREP vector served as a negative control. Transformation reactions were plated on an average of 10 agar plates (ø 140 mm) containing 50 μg/mL kanamycin and 34 μg/mL chloramphenicol such that each plate contained ∼1, 000 colonies, equivalent to a screening capacity of ∼10,000 clones. Colonies were blotted onto nitrocellulose membranes (BioTrace NT 0.2 μm, PALL Corp), and transferred facing up onto a second LB agar plate containing 1 mM IPTG, 200 μM ZnCl_2_ (to ensure availability of Zn^2+^ ions necessary for enzymatic activity) and 0.2% (w/v) arabinose for chaperone overexpression. The original agar plates were placed at 37 °C for colonies to re-grow, while the membrane-containing plate was left overnight at room temperature for expression. The next day, membranes were placed into an empty petri dish and cells were lysed by three freeze/thaw cycles at -20 °C and 37 °C. For the activity assay, 25 mL of 0.5% agarose dissolved in 50 mM Tris-HCl buffer, 100mM NaCl, pH 7.5 containing 200 μM ZnCl_2_ and 200 μM 2NH with 1mM Fast Red were poured onto the nitrocellulose membranes. Red colour developed within 10-60 minutes. Positive clones were then picked from the original agar plate (see arylesterase assay in cell lysate for the next step).

#### Arylesterase assay in cell lysate

*Directed evolution screening* - Active clones identified in the colony screening were re-grown overnight at 30 °C in 4-6 deep 96-well plates containing 200 μL Lysogeny broth (LB) supplemented with 50 μg /mL kanamycin and 34 μg/mL chloramphenicol for pGro7 selection. This led to a focused library of ∼360-560 pre-selected variants. Subsequently, 25 μL of overnight cultures were transferred to deep well plates containing 425 μL LB per well with kanamycin, chloramphenicol, 200 μM ZnCl_2_ and 0.2 % (w/v) of arabinose for chaperone co-expression. Cells were grown for ∼2H at 37 °C until the OD reached ∼0.6. Expression of PTE variants was induced with 1 mM IPTG and cultures were incubated for 2 hours at 30°C. Cells were spun down at 4 °C at maximum speed (3320 g) for 10 minutes and the supernatant drained. Pellets were frozen for > 1H at -80 °C and subsequently lysed by addition of 200 μL of 50 mM Tris-HCl buffer, 100mM NaCl, pH 7.5 supplemented with 0.1% (w/v) Triton-X100, 200 μM ZnCl_2_, 100 μg/mL lysozyme and 1 μL of benzonase (Novagen, 25 U/μL) per 100 mL of lysis buffer. After 30 minutes of lysis, cell debris were spun down at 4 °C at 3320 g for 30 minutes. Depending on the activity level of the library, the clarified lysate was diluted prior to the activity assay to obtain a good signal in the initial linear phase of the reaction. Reactions were performed in 96-well plates (Corning Costar) containing 200 μL per well (20 μL lysate + 180 μL of the substrate (200 μM of 2NH + 1mM of Fast Red) previously dissolved in the same lysis buffer. 2NH hydrolysis was monitored at 500 nm via complex formation with Fast Red. ACP (acyl phosphatase) cloned into the same pET-27-STREP vector served as a negative control for 2NH activity measurements. Improvements ≥1.3-fold relative to the previous round were considered significant. The best clones were picked, re-grown, and re-assayed in triplicate. The observed initial rate values were normalized to cell density (determined by absorption at 600 nm) and the average values were determined. Around 10 improved variants were sequenced after each round. A description of each directed evolution round including selection criteria, the mutations found in each sequenced variant, and mention of the variants chosen as templates for the next library generation, can be found in the SI. *PTE mutants assay* - Single point PTE mutants were assayed as described above, except for the overnight cultures being inoculated directly from glycerol stocks in triplicates in 2-3 independent experiments. Saturation mutagenesis libraries were first plated on agar plates where ∼85 colonies were randomly picked for overnight cultures, prior to lysate screening as described above.

#### Protein purification

*STEP-tag purification* - Selected evolved PTE variants were purified using strep-tactin affinity columns according to the manufacturer’s instructions (IBA BioTAGnology, Germany). Briefly, pET-Strep-PTE plasmids were transformed into *E. coli* BL21 (DE3) (without pGro7 chaperone expression for higher purity) and a single colony was used to grow a 2 mL LB culture overnight at 37 °C. Cells were inoculated into Overnight Express Instant TB medium (Novagen) containing 1% (v/v) glycerol, 50 µg/mL kanamycin, 200 μM ZnCl_2,_ and STREP-tagged proteins were expressed at 30°C for 8H at 250 rpm, followed by 12H at 16°C. Cells were harvested by centrifugation and pellets were frozen overnight at -80°C. Pellets were resuspended with 20 mL Lysis buffer containing a 1:1 mixture of B-PER Protein Extraction Reagent (Thermo Scientific): 50 mM Tris-HCl buffer, 100mM NaCl, pH 7.5 supplemented with 200 μM ZnCl_2_, 100 μg/mL lysozyme and ∼0.5 μL of benzonase (Novagen). After a 1H-incubation on ice, cell debris were removed by centrifugation and the clarified lysate passed through a 45 μm filter before loading onto a Strep-Tactin Superflow High-capacity column (IBA, CV=2.5 mL). Strep-PTE variants were eluted with Elution buffer (50 mM Tris-HCl, 100 mM NaCl, pH 7.5 with 200 μM ZnCl_2_ and 2.5 mM desthiobiotin) according to the manufacturer’s instructions (IBA). Approx. 3mL of PTE-containing pure fractions (determined by SDS-Page and 2NH activity measurement) were subsequently pooled, loaded onto an Econo-Pac 10DG desalting column (Bio-Rad) and eluted with the same Elution Buffer without desthiobiotin. When needed, PTE variants were concentrated to 0.5-2 mg/mL using a 10K MWCO spin concentrator (Microsep Advance centrifugal device, PALL Corp), their concentrations were measured in triplicate using the Pierce BCA Protein assay kit (Thermo Scientific) and they were stored at 4 °C.

##### Purification of untagged variants for protein crystallization

PTE variants were cloned into pET32-trx plasmid (ampicillin) without Strep-tag using FastDigest NcoI and HindIII as described in the cloning section, transformed into *E. coli* BL21 (DE3), and grown for 72 hr at 30 °C in TB medium containing 100 μg/ml ampicillin and 500 μM ZnCl_2_. Cells were harvested by centrifugation at 3, 320×g and 4 °C for 10 min, resuspended in 20 mM Tris-HCl, pH 8 containing 100 μM ZnCl_2,_ and lysed by sonication (OMNI Sonic Ruptor 400, Thermo Scientific, Waltham, MA, United States, 50% pulse, 50% amplitude). Cell debris were removed by centrifugation at 30,000×g and 4 °C for 60 min and the lysate was filtered through 45 μm filters (Millipore). The lysate was loaded onto a Fractogel DEAE anion exchange column (Merck). PTE elutes in the flow-through, as well as in the early wash fractions. Fractions deemed to be >95% pure and active were pooled. Protein variants were concentrated to 12 mg/mL and stored at 4 °C.

#### Enzyme kinetics

Initial velocities (v_0_) were determined in duplicate, at 16 different substrates concentrations (0–2 mM) in 50 mM Tris-HCl, 100 mM NaCl, pH 7.5, 200 μM ZnCl_2_ and 0.1% Triton X-100, at 25 °C. Hydrolysis rates were monitored using a microplate reader (Synergy H1, Biotek) by following product formation at 500 nm for 2NH (in the presence of FAST-Red, at a ratio of 1:5). Steady-state kinetic parameters were derived by fitting the experimental data to the Michaelis-Menten equation: v_0_/[E] = k_cat_ · [S]/(K_M_ + [S]), where v_0_ is the initial rate, [E] is the enzyme concentration, and [S] is the substrate concentration. In some cases, we observed substrate inhibition or could not reach full saturation, in which case we used a linear fit: v_0_/[E] = (k_cat_ /K_M_) · [S] or a substrate inhibition fit: v_0_/[E] = k_cat_ · [S]/(K_M_ + [S] · (1 + [S]/ K_i_)) where K_i_ represents the inhibition constant. The data were corrected for the buffer-catalysed background reaction measured under the same conditions without enzyme. All steady-state parameters and detailed procedures can be found in SI Materials and Methods.

#### Linear modeling of genetic effects

##### Definition of genetic encoding system

To quantify the genetic and environmental determinants of enzyme activity, we used an approach similar to that previously developed ^30,31^. We constructed regression models that explain the enzyme activity (*E.A.*) as a function of the genetic states at three variable amino acid positions (254, 233, and 271), and varying across three different states: (*i*) the ancestral state, and the unique derived states found in the (*ii*) *Ser* and (*iii*) *Arg* trajectories, respectively. The genetic variation in the protein was defined in the linear models using one-dimensional variables for each mutation, going from ancestral to derived state, calculated independently along each trajectory, and then separately when combining mutations from different trajectories.

##### First-order linear models

We constructed our first-order model by regressing the enzyme activity of each genotype on dependent variables that reflect the individual first-order identities at each genetic position. For example, the linear model for position 254 is expressed as:

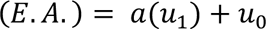

where *a* is the effect coefficient of moving +1 in that dimension, *u_1_* is the coordinate representing the genotype (*i*.*e*., 0 for ancestral leucine, +1 for derived arginine), and *u*_0_ is the y-intercept for the model. The linear coefficients for each model were computed using ordinary least squares (OLS) regression with the open-source statistical package R (http://www.r-project.org/). The coefficient *a* indicates the deviation of the derived genetic state from the ancestral. To determine how well all possible first-order effects of mutations along a specific trajectory in the protein predict variation in *E.A.*, we constructed the following linear model that included all first-order protein coefficients:

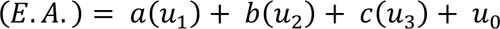

where *u*_2_, *u*_3_, are the coordinates representing the genotype at positions 233 and 271, respectively. We then computed the R^2^ for this first-order model.

##### Linear models with second-order genetic epistasis

As second-order models can be computed and compared simply, we added an interaction factor for each pairwise combination (*e.g.*, for 254×233) and compared the R^2^ of that model to that of the first-order model. We then computed a likelihood ratio test between the more complex and the simpler model to assess whether epistasis was statistically significant in each case.

#### Evolutionary pathway assessment

To model the evolutionary pathways that would have been available to the enzyme under a model of strong direction selection (*i*.*e*., classic Darwinian adaptation), we assessed for each possible mutation whether a “step” toward the derived state was available (*i.e.*, by increasing enzyme activity) or not. The experimental trajectories we determined are each highlighted on the fitness landscape as vertical lines connecting genotype nodes, while the availability of a step from the sub-optimal trajectory (*i.e.,* the *Ser* trajectory) to the optimal trajectory (*i.e.,* the *Arg* trajectory) is shown as red (not available) or blue (available) lines.

#### Crystallization of PTE variants

Purified untagged PTE variants were concentrated to 12 mg/mL in 20 mM HEPES, 50 mM NaCl, 100 μM ZnCl2, pH 8. Crystals were grown at 4 °C in EasyXtal-15-Well Tools plates (Qiagen), using hanging-drop vapour diffusion. Drops consisting of 2 μL of reservoir solution and 1 μL of PTE were equilibrated over 500 μL of reservoir solution. Reservoir solutions contained 100 mM sodium cacodylate (pH 6.3 - 6.7), distilled deionized water, and 10-50% (w/v) 2-methyl-2,4-pentanediol (MPD). Crystals formed in initial screens were manually pulverized and used for microseeding. Six serial dilutions of the pulverized crystals were made, from 1/10 to 1/1,000,000 in factors of 10. New crystal screens were set up with varying MPD concentrations from 10% to 20%. Each well contained 1 μL of protein, 1.5 μL of reservoir solution, and 0.5 μL of the microseed solution. To obtain crystal structures in the same space group and crystallization condition, serial microseeding was used. Co-crystallization of PTE variants with the transition state analogue HLN was achieved by adding a 4 times molar excess of HLN to crystallization wells. Diffraction-quality crystals typically formed within a week.

#### X-ray diffraction data collection and refinement

For data collection, crystals were soaked in cryobuffer (100 mM sodium cacodylate, pH 6.5, 40% MPD) for 2 minutes before flash-cooling to 100 K in a stream of nitrogen gas. PTE diffraction data were collected at the Australian Synchrotron (beamlines MX1^32^ and MX2 ^33^) at a wavelength of 0.9537 Å. The X-ray diffraction data were indexed and integrated using XDS ^34^ and merged using AIMLESS as implemented within the CCP4 program suite ^35^. Data resolution cut-offs were chosen using half-dataset correlation coefficients, as described by Karplus and Deiderichs ^36^. The structures of PTE variants were solved by molecular replacement (MOLREP) ^37^ using the PTE^WT^ structure (PDB 4PCP). Refinement was performed using REFMAC and phenix.refine, and manual rebuilding was performed in Coot ^38^. Automated fitting of multiple conformations was carried out with the QFit web-server ^39^, with manual inspection and removal of conformations that were not present in good (at least 1.0 σ) electron density. Network analysis was performed using RING 2.0 ^40^, and results were visualized with Cytoscape ^41^. Cα RMSD calculations were performed using FATCAT ^42^.

#### B-factor and B’-factor analysis

Isotropic alpha-carbon B-factors were extracted from each structure. For residues with multiple conformers modelled, conformer B was omitted from the analysis. Baseline-normalization was performed using a modified z-transformation method ^43^ to obtain B’-factors for each alpha carbon (B’i): Regions of high variance (*e.g.*, loops and termini: residues 0-150, 170-178, 20-210, 230-239, 255-280, 300-450) were omitted when calculating the mean baseline B-factor 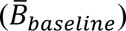 and standard deviation of the B-factor (σ_*baseline*_) of each chain prior to calculating *B*′(*i*) (equation 1). Scripts for analysis and plot preparation, as well as full-resolution figures, are available on GitHub (https://github.com/jkaczmarski/PTE_PAPER_2023).

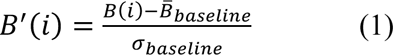

#### Ensemble Refinement

Phenix.ensemble_refinement (PHENIX version 1.18.2) ^44^ was used to generate time-averaged ensembles of S1, S2, S3, S5, S6, S7 and S8 (structures from this study), and R0, R1, R2, R6, R8, R18 and R22 (from ref. ^18^). Structures and structure factors were obtained from the wwPDB and re-refined using phenix.refine. For each structure in both Ser and Arg trajectories, phenix.readyset was used to add hydrogen atoms and generate .cif files for the ligands. Each monomer present in the crystallographic dimer was assigned to a unique translation-liberation-screw (TLS) group. Harmonic restraints were added to any ligands present. The ensemble refinement parameters were optimized by testing various TLS fitting procedures (*p*_*TLS*_= 0.6, 0.8, 0.9, 1.0) followed by optimization of the X-ray weight (by changing the simulation temperature bath, T_bath_ = 290 K, 295 K, and 297.5 K) and the relaxation time of the time-averaged restrained (T_x_ = 0.3, 0.6, 1). Finally, nine additional random seed repeats were performed using the parameters that led to the ensemble with the lowest R_free_. Alpha-carbon RMSF values were calculated using the get_rmsf tool in GROMACS (version 2018.3); analysis was performed on the five ensembles with the lowest R_free_ values, following averaging of values across the two chains in the crystallographic dimer. Scripts used for analysis, plot preparation and PyMOL figures, as well as full resolution images, are available on GitHub (https://github.com/jkaczmarski/PTE_PAPER_2023).

#### Molecular dynamics simulation

Molecular Dynamics simulations of PTE variants R0, R1, R18, S1, S5, and S8 were performed in explicit water using AMBER20 package ^45^. The protein was described with the AMBER-ff14SB force field ^46^, the two active-site zinc ions coordination sphere was represented with a bonded model generated with MCPB.py ^47^ and GAFF parameters ^48^, and water molecules with the TIP3P force-field ^49^. Each variant was solvated in a cubic box with a 10 Å buffer of TIP3P water molecules and was neutralized by addition of explicit sodium and chloride counterions (Na^+^ or Cl^-^). Next, a two-stage geometry optimization approach was performed. First, a short minimization of the water molecules positions, with positional restraints on solute by a harmonic potential with a force constant of 500 kcal mol^-1^ Å^-2^ was done. The second stage was an unrestrained minimization of all the atoms in the simulation cell. Then, the systems were progressively heated using six 50 ps steps, incrementing the temperature 50 K each step (0-300 K) under constant-volume, periodic-boundary conditions and the particle-mesh Ewald approach to introduce long-range electrostatic effects ^50^. For these steps, a 10 Å cut-off was applied to Lennard-Jones and electrostatic interactions. Bonds involving hydrogen were constrained with the SHAKE algorithm. Harmonic restraints of 10 kcal mol^-1^ were applied to the solute, and the Langevin equilibration scheme is used to control and equalize the temperature. The time step was kept at 2 fs during the heating stages, allowing potential inhomogeneities to self adjust. Each system was then equilibrated for 2 ns with a 2 fs time step at a constant pressure of 1 atm to relax the density of the system. After the systems were equilibrated in the NPT ensemble, three replicas of 500 ns MD simulations for each variant (R0, R1, R18, S1, S5, and S8) were performed under the NVT ensemble and periodic-boundary conditions using the Galatea cluster at the University of Girona.

#### Shortest Path Map analysis

The MD simulations were further analyzed by calculating the inter-residue mean distance and correlation matrices for SPM construction. These matrices were then used to construct a first complex graph, in which all residues that present a mean distance value of less than 6 Å are connected with a line weighted according to their correlation value (d_ij_=-log |C_ij_|). This first graph is further processed to identify the shortest path lengths, *i.e.*, the ones playing the most important role in the conformational dynamics of the enzyme ^19,20^. The final SPM graph is then represented on top of the tridimensional structure.

#### Data deposition

The apo structures of PTE-S1 (PDB 5W6B), PTE-S2 (PDB 5WCQ), PTE-S3 (PDB 5WCW), PTE-S5 (PDB 5WIZ), PTE-S5+ΔL7 (PDB 6BHL), PTE-S5+254R (PDB 5WJ0), PTE-S6 (PDB 5WCP), PTE-S7 (PDB 5WMS), PTE-S8 (PDB 6AML), and of PTE-S5 in complex with a 2NH analogue (PDB 6B2F), have been deposited in the RCSB Protein Data Bank, www.rcsb.org.

## Authors contribution

C.M.M., E.C.C. J.A.K, FF, A.R.,M.S,, D.A, and N.S performed experiments and analyzed results. F.F. and A.R performed MD simulations and SPM analysis. C.J.J, J.A.K and M.S. performed ensemble refinement and data analysis. E.C.C purified, crystallized, solved and analyzed the structure of all variants in this study. D.A. designed custom codes and performed the statistical analyses. C.M.M performed all other experiments, C.M.M., N.T., C.J.J. and S.O conceived the project, designed experiments, analyzed results and wrote the manuscript. All authors contributed to writing this article.

## Acknowledgments

We thank the members of the Tokuriki, Jackson and Osuna laboratories for their invaluable comments. We would like to thank our friend Dan S. Tawfik for his benevolent contribution, stimulating discussions and insightful advice about this particular project. N.T. received the support of the Natural Sciences and Engineering Research Council of Canada (NSERC) Discovery Grant (RGPIN 2017-04909). N.T. and S.O. are supported by the Human Frontier Science Program (HFSP) Research Grant (RGP0054/2020). C.J.J is supported in part by the Australian Government through the Australian Research Council Centres of Excellence funding scheme (projects CE200100029 & CE200100012). This research was undertaken on the MX1 and MX2 beamlines at the Australian Synchrotron, Victoria, Australia. S.O. and F.F. thank Generalitat de Catalunya AGAUR for 2021SGR00487. F.F. is supported by the Spanish MINECO for RYC2020-029552-I and PID2022-141676NB-I00. S.O. received the support of the European Research Council (ERC) under the European Union’s Horizon 2020 research and innovation program (ERC-2015-StG-679001, ERC-2022-POC-101112805, and ERC-2022-CoG-101088032) and the Spanish MICIN for grant projects PID2021-129034NB-I00 and PDC2022-133950-I00.

